# Detecting natural hybridization among Indian major carps through advance morphometric approach

**DOI:** 10.1101/553941

**Authors:** Arvind Kumar Dwivedi

**Author notes:** **Correspondence**: (AKD).

## Abstract

Interspecific natural hybridization is an indicator of altered ecosystem. Habitat destruction increases competition with fish species in close proximity for spawning habitat with overlapping reproductive activities, thereby causing natural hybridization. This study is first investigation on detecting hybrids among Indian major carps (*Labeo rohita, Cirrhinus mrigala* and *Gibelion catla*) from the Ganga River by applying a cost-effective method, “Geometric Morphometrics”. The relative warps (RW), principal component analysis (PCA) and canonical variate analysis (CVA) were employed on superimposed images to determine morphometric variations. Deformation grid of RW showed that *G. catla* and hybrid specimens have a deeper whereas *L. rohita* and *C. mrigala* specimens have slender body profile. The PCA showed separation among specimens of four groups (three species and one hybrid) with slight overlap between *G. catla* and hybrid. CVA extracted Mahalanobis and Procrustes distances among four non-overlapping groups found to be highly significant (P < 0.0001) with hybrid specimens lying between position of *L. rohita* and *G. catla* specimens in close proximity to *G. catla*, suggesting that hybrids are product of crossing between *L. rohita* and *G. catla.* The CVA separated four groups with 100.00% classification, indicating that all the three species and hybrid were clearly distinct from each other. In this study, all the four specimens of hybrid were caught from upstream of four barrages (Bijnor, Narora, Kanpur and Farakka) commissioned along the middle and lower stretch of the Ganga River. This suggests that, barrages obstruct upward movement of fish population and impact on reproductive activities, thereby increases possibilities of natural hybridization among these species.

## Introduction

River Ganga, India’s National River is home to a wide variety of relic species, is fragmented by dams and barrages constructed along its entire stretch. Over the last few decades, wild capture fishery in Ganga River appears to be declined [1] and changes in fish distribution, phenotypic traits and biological characteristics have also been reported [2]. Indian major carps (Order-Cypriniformes, Family-Cyprinidae) are economically important warm-water teleost, inhabitant of Indo-Gangetic riverine system spread across northern and central India, and the rivers of Pakistan, Bangladesh, Nepal and Myanmar [3, 4]. In the Ganga River, Indian major carps (IMCs) are distributed all along the middle (Bijnor to Patna) and lower stretch (Patna to Hooghly) [5]. They have been effectively transplanted out of its natural range within India and parts of Asia as well as Europe [6, 7]. The major source of seed in India is contributed from natural resources. Indian major carps (*Labeo rohita, Cirrhinus mrigala* and *Gibelion catla*) contribute nearly 4% to the global aquaculture [8]. Hybridization among these Indian major carps is most common in nature, especially between *L. rohita* and *G. catla* [9-12].

Cyprinid fishes are prone to interspecific hybridization in wild [13]. Several causes have been suggested to account for the occurrence of hybrids in nature. Hubbs [14] holds the view that hybridization in nature is facilitated when there is scarcity of one species and dominance of an allied species in close proximity. This phenomenon is particularly common when populations invade new environment and overlap temporally and spatially in spawning activities [15-18]. This tendency is further enhanced by anthropogenic factors, such as habitat destruction, which increases competition for spawning habitat [19, 20]. Many studies have stressed that interspecific hybrids exhibit significant reductions in viability, fertility, and fitness [21-22].

Misidentification of pure and hybrid fingerlings of Indian major carps from wild may lead to hybridization during aquaculture practices subsequently deteriorating the quality of seed produced [23-25]. During fish diversity assessment, misidentification of hybrids may lead to taxonomic ambiguities. Few studies have been conducted in recent past to differentiate hybrids from pure fingerlings employing various tools such as phenotypic traits, allozymes and mitochondrial DNA markers [25-27]. However, the phenotypic traits described in these studies are more or less complex and molecular techniques used are also very costly. Therefore, the present scenario demands a cost effective, reliable and user friendly method to detect hybrids among Indian major carps.

It is also obvious that over the past few years, science of taxonomy has been suffering from dwindling number of experts [28]. Moreover, the pace of traditional taxonomy is also very slow. Taxonomists carefully count meristic characters, measure multiple morphometric variables of body shape and examine external body features to discriminate Indian major carps, still there is a risk of misidentification during visual assessment [29]. However, in the recent past, the pace of data gathering and analysis in taxonomy has been greatly increased by the development of information technology [30]. A family of software tools have been designed for gathering and analyzing data on shape variation from images of specimens [31]. “Geometric Morphometric” (GM) method is currently considered to be the most rigorous, cost effective and user friendly morphometric technique in capturing information about the shape of an organism from digital images with more powerful statistical analyses for species differentiation [18, 32-34].

Till date, no study has examined shape variation among Indian major carps and detecting their hybrids using “Geometric Morphometrics” in nature. During fish diversity assessment of Ganga River, four individuals were encountered, which were suspected to be the hybrids of *L. rohita* and *G. catla*, having scale colour and pattern of *L. rohita* and body profile especially head region and body depth profile of *G. catla*. To resolve the taxonomic ambiguity, this study was conducted with the aim to evaluate morphometric variation among Indian major carps and detecting their hybrids using “Geometric Morphometric” method to provide new insights into features of shape change. By examining the external morphology and distribution range of the Indian major carps and their hybrids in the Ganga River, shape variations of the latter from parental species were explored along with anthropogenic factors responsible for their successful adaptation.

## Materials and Methods

### Study areas and sample collection

A total of 96 specimens of Indian major carps including 4 individuals of hybrids were collected with the help of local fishermen using different fish gears (gill net, drag net and cast net) from ten locations along the middle and lower stretch of Ganga River between May 2017 and April 2018. The sample size and relative information for all the sampling sites are presented in Table 1 and Fig 1.

**Table 1:**
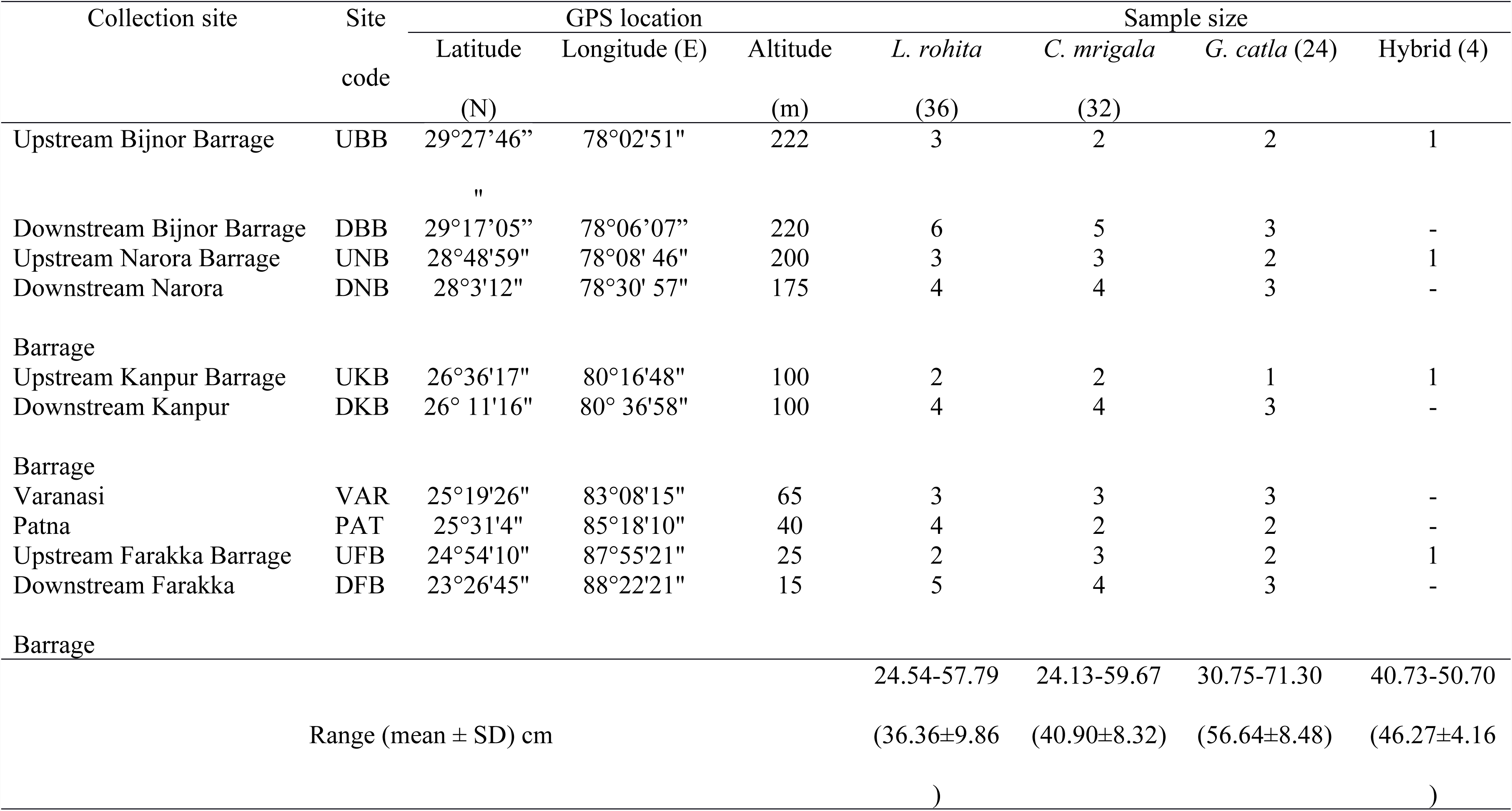
Collection localities, sample size and size statistics (based on the Standard Length) of Indian major carps in Ganga River

**Fig. 1.**
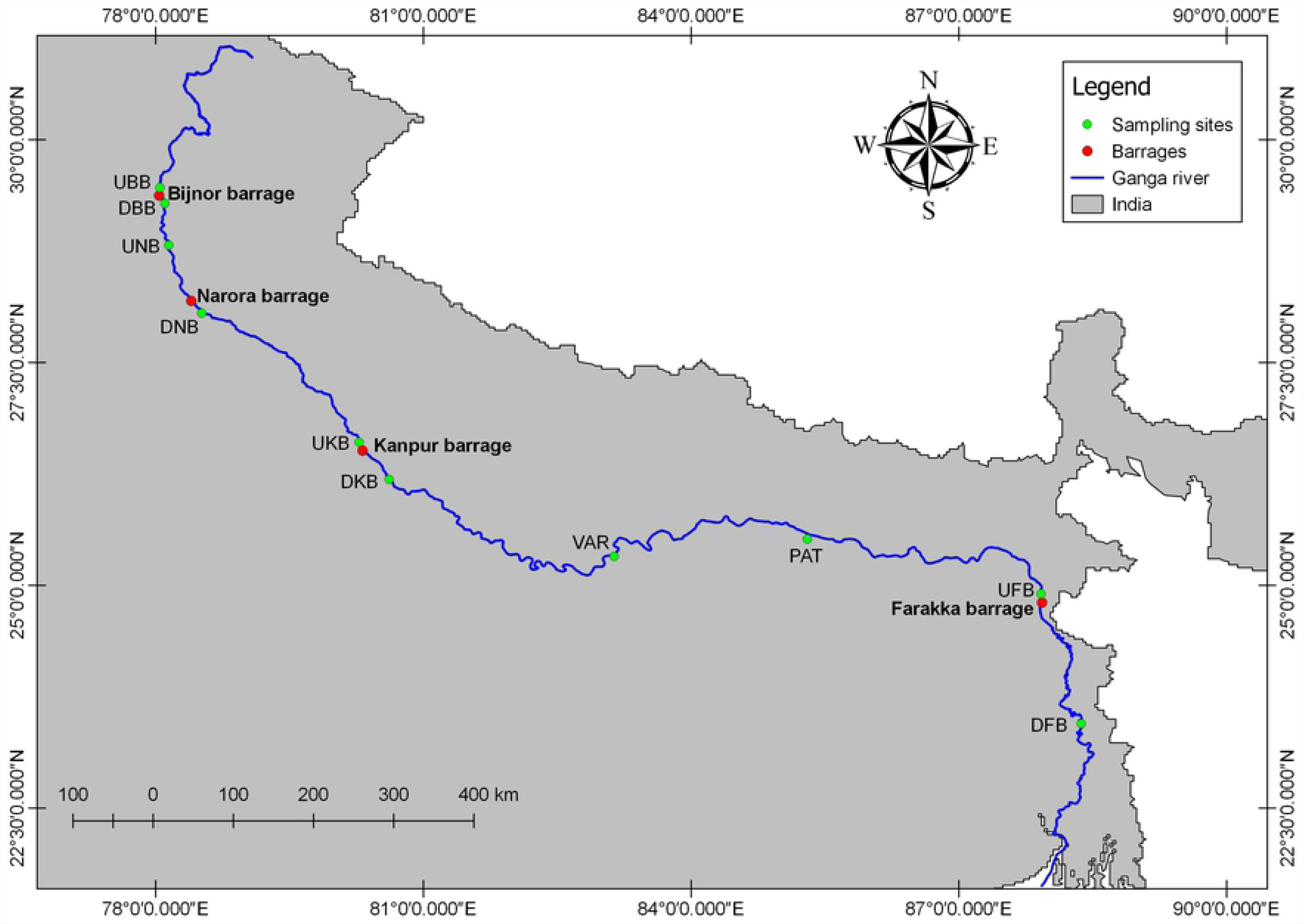
Map of Ganga River showing collection area of Indian major carps and hybrids

### Ethics statement

As per IUCN status, Indian major carps are under Lease Concern (LC) category. Fish samples were obtained from wild and kept live in water tanks. Sites from where fishes were collected fell outside Protected Areas (PAs) and therefore no permits were required from State Forest and Wildlife Department. After taking photographs, fishes were released back to the river system. No fish samples were sacrificed or anesthetised. Therefore, ethics committee approval not required.

### Geometric morphometric analysis

Sampled specimens were uniformly placed on laminated graph sheets, body posture and fins were teased into a natural position. Each individual was labelled with a specific code for identification. A Cyber shot DSC-W300 digital camera (Sony, Japan) was used to capture the digital lateral image of the left side of each individual.

TPSUTIL, version 1.52 was used to build a tps file from the photographs [35]. On each specimen, 14 homologous landmarks were digitized using the software TPSDIG version 2.16 [36] (Fig 2). Scale factor using set scale option was performed on each specimen. 2D X-Y coordinate data for 14 landmarks of all the 96 specimens were generated using software ImageJ version 1.50i (http://imagej.nih.gov/ij/) and saved in tps format.

**Fig. 2.**
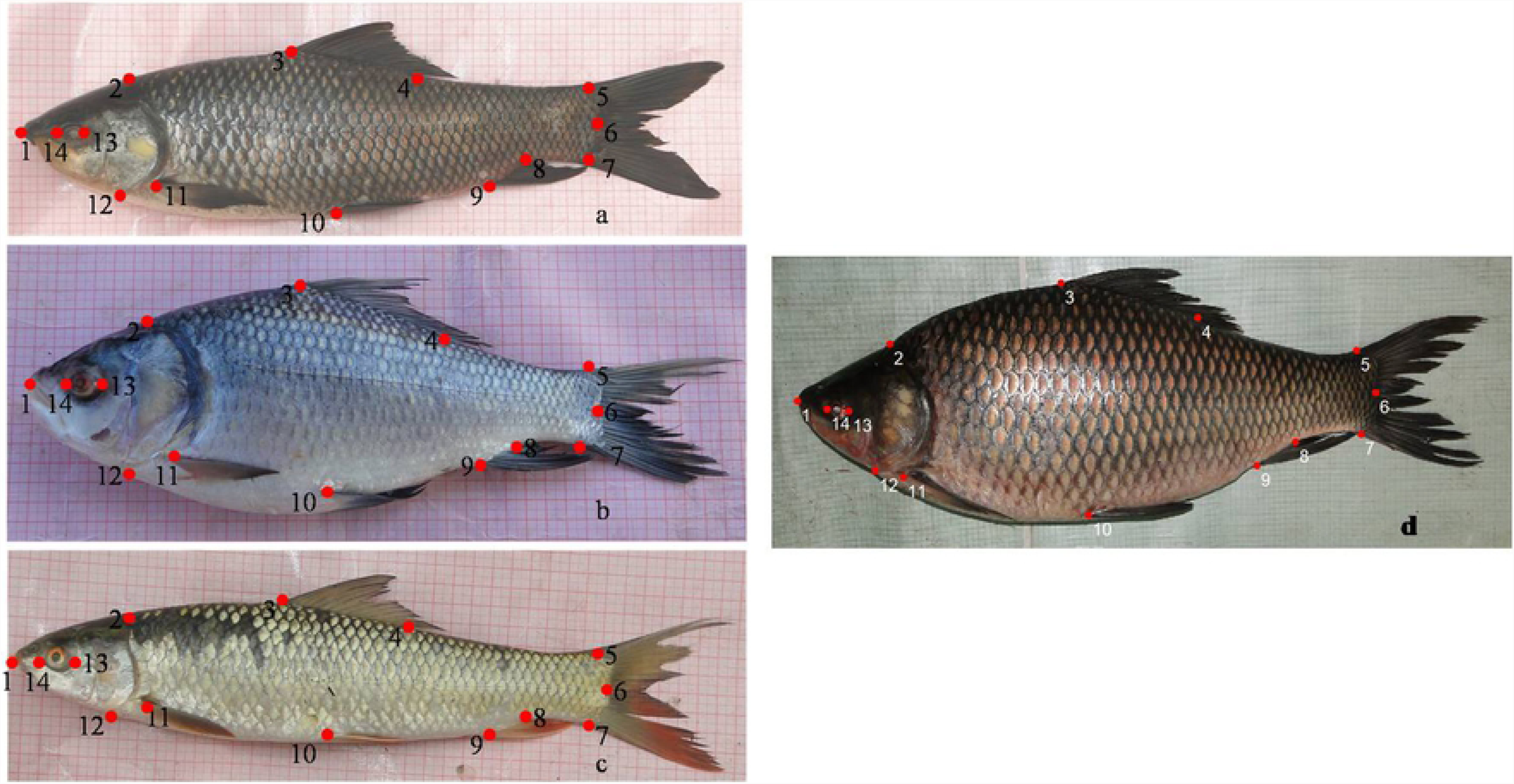
Location of 14 landmarks on Indian major carps a. *L. rohita*, b. *G. catla*, c *C. mrigala*. Landmarks refer to 1. anterior tip of snout at upper jaw 2. most posterior aspect of neurocranium (beginning of scaled nape) 3. origin of dorsal fin 4. end of dorsal fin 5. anterior attachment of dorsal membrane from caudal fin 6. posterior end of vertebral column 7. anterior attachment of ventral membrane from caudal fin 8. end of anal fin 9. origin of anal fin 10. insertion of pelvic fin 11. insertion of pectoral fin 12. ventral edge of the operculum 13. posterior end of eye 14. anterior end of eye

Landmarks were converted to shape coordinates by Procrustes superimposition [37], standardizing each specimen to unit centroid size, or an estimate of overall body size [38]. The pattern of covariation between shape and size as well as shape and sex was analysed using Partial Least Square (PLS) analysis [39, 40]. Relative warp (RW) analysis was performed using TPSRELW [41] to synthesize and visualize the morphological variations. Wireframes connecting the digitized landmarks on a shape were generated to visualize and interpret shape changes between the two seasonal migrants. Principal Components Analysis (PCA) and Canonical Variate Analysis (CVA) were performed to assess the shape variation between the two seasonal migrants. Principal components analysis was applied to outline groups of samples and to identify influential variables [42]. Canonical variate analysis is a method of finding the set of axes that allows for the greatest possible ability to discriminate between two or more groups [43]. All statistical analyses were carried out with the software MorphoJ, version 1.06d [44].

## Results

Shape coordinates were successfully superimposed by Procustes analysis (Procrustes sums of squares: 0.363 and Tangent sums of squares: 0.361). PLS revealed a non-significant covariance between shape and size (R = 0.54; P > 0.001). In this case, plot shows overlapping among groups (Fig 3).

**Fig. 3.**
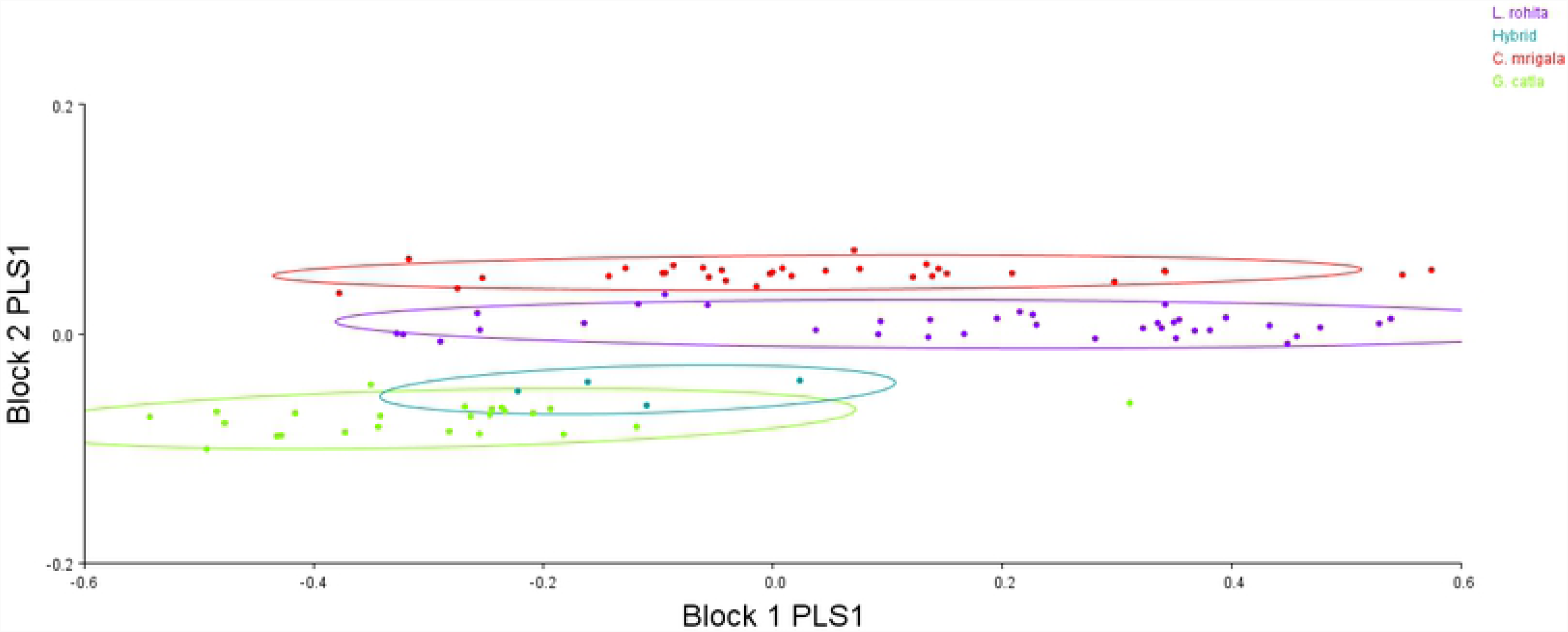
Scatter plot of the partial least square analysis in Indian major carps and hybrids computed on size (Block 1 PLS1) and shape (Block 2 PLS1) variables

RW analysis showed deformation in shape from the reference that corresponds to selected positions in the ordination. As shown by the deformation grid, *L. rohita* and *C. mrigala* have slender and *G. catla* and hybrid have deeper body profile. Deformed wireframe was drawn on the shape of three species and hybrid to interpret shape changes which supports the relative warp analysis (Fig 4).

**Fig. 4.**
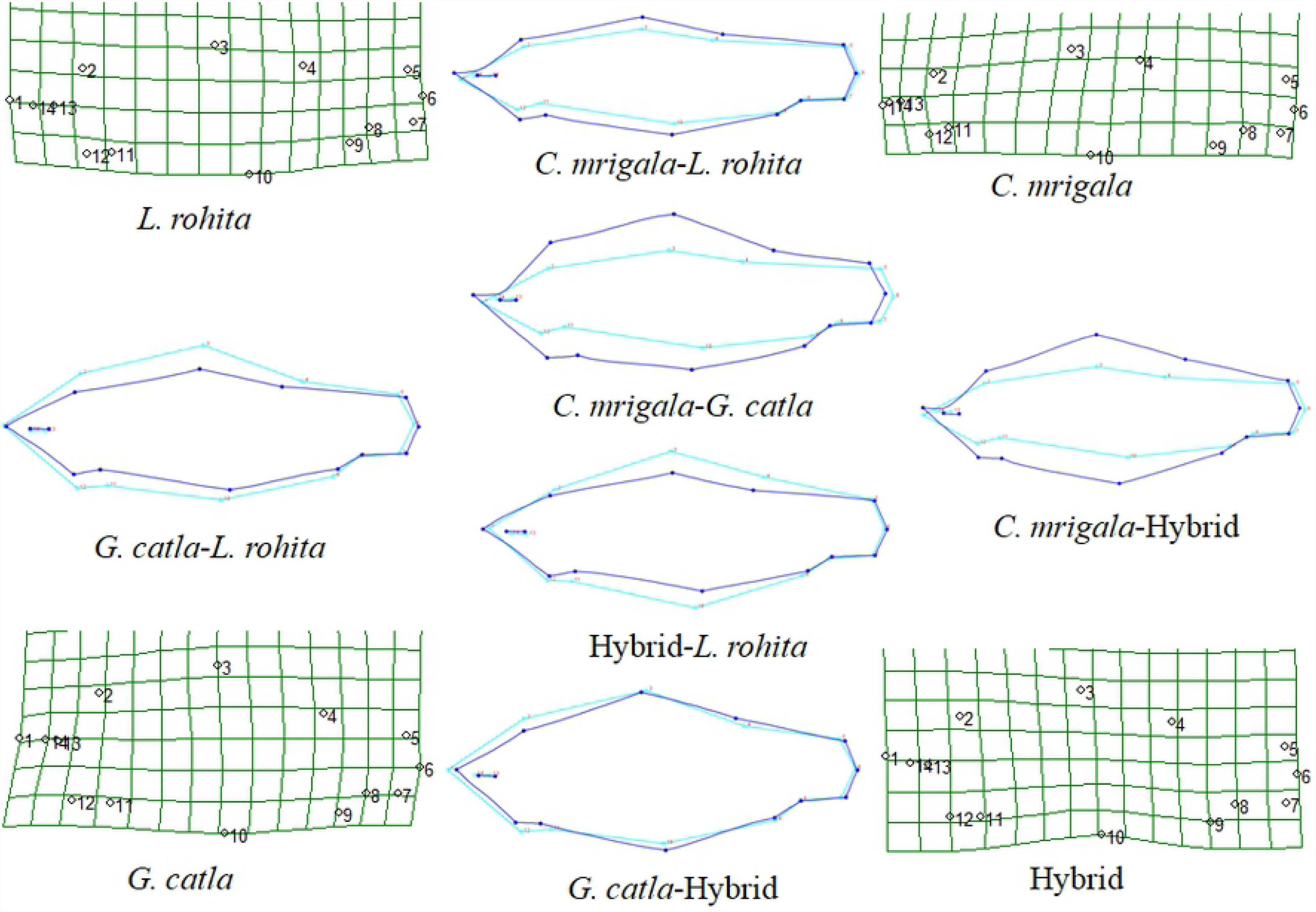
Deformation grid of relative warps and wireframes between two groups in Indian major carps and hybrids

The PCA extracted 24 components with 100.00% variance (Fig 5). The first two principal components (PC) account for 79.66% of the total variance (74.32% for PC1, 5.34% for PC2). Separation among four groups with slight overlap between *G. catla* and hybrid was clearly evident in the PCA plot of the first against the second PC axes (Fig 6). CVA based upon 14 landmarks showed four groups with no overlaps among groups with position of hybrid specimens lying between position of *L. rohita* and *G. catla* specimens in close proximity to *G. catla* (Fig 7). CVA plot suggested that hybrids are resultant due to crossing between *L. rohita* and *G. catla.* CVA extracted three CVs accounting for 100.00% variation (92.47% for CV1, 5.86% for CV2). CVA extracted Mahalanobis and Procrustes distances among four groups found to be highly significant (P < 0.0001), indicating that all the four groups were clearly distinct from each other (Table 2, 3). Classification results of CVA indicated that all the specimens of each group were assigned to their respective group with no misclassification rate (Table 4).

**Table 2:** Mahalanobis distances among groups (upper diagonal) and *p* value (lower diagonal) of canonical variate analysis

| Groups | *L. rohita* | *C. mrigala* | *G. catla* | Hybrid |
| --- | --- | --- | --- | --- |
| *L. rohita* |  | 8.491 | 13.580 | 10.210 |
| *C. mrigala* | <0.0001 |  | 20.350 | 16.947 |
| *G. catla* | <0.0001 | <0.0001 |  | 7.443 |
| Hybrid | <0.0001 | <0.0001 | <0.0001 |  |

**Table 3:** Procrustes distances among groups (upper diagonal) and *p* value (lower diagonal) of canonical variate analysis

| Groups | *L. rohita* | *C. mrigala* | *G. catla* | Hybrid |
| --- | --- | --- | --- | --- |
| *L. rohita* |  | 0.055 | 0.085 | 0.067 |
| *C. mrigala* | <0.0001 |  | 0.134 | 0.114 |
| *G. catla* | <0.0001 | <0.0001 |  | 0.050 |
| Hybrid | <0.0001 | <0.0001 | <0.0001 |  |

**Table 4:** Classification of individuals/percentage in respective group (in diagonal), misclassification of individuals (upper diagonal) and their percentage (lower diagonal)

| Groups | *L. rohita* | *C. mrigala* | *G. catla* | Hybrid |
| --- | --- | --- | --- | --- |
| *L. rohita* | 36/100.00 | 0 | 0 | 0 |
| *C. mrigala* | 0.00 | 32/100.00 | 0 | 0 |
| *G. catla* | 0.00 | 0.00 | 24/100.00 | 0 |
| Hybrid | 0.00 | 0.00 | 0.00 | 4/100.00 |

**Fig. 5.**
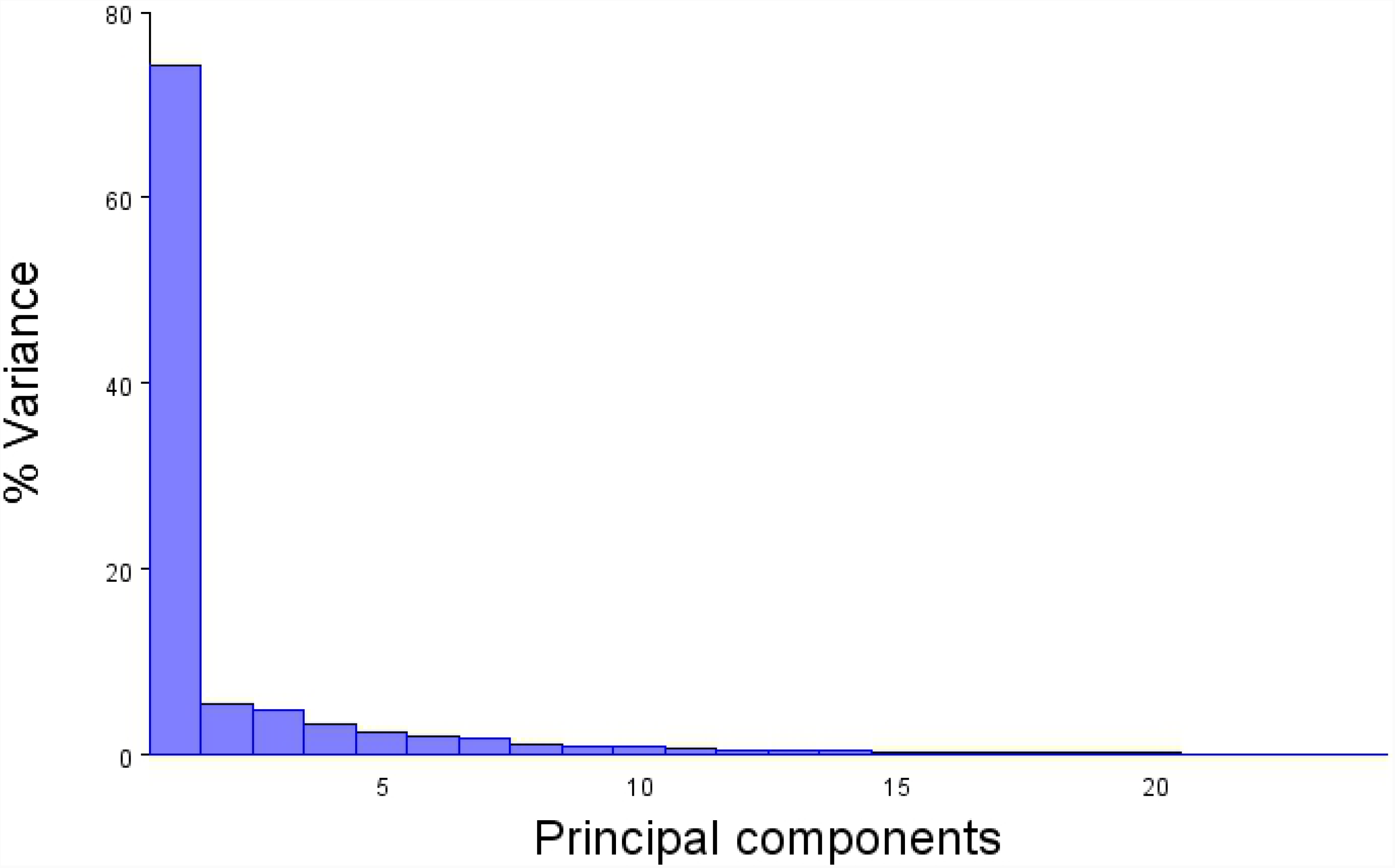
Scree plot of percentage variance and principal components for Indian major carps and hybrids

**Fig. 6.**
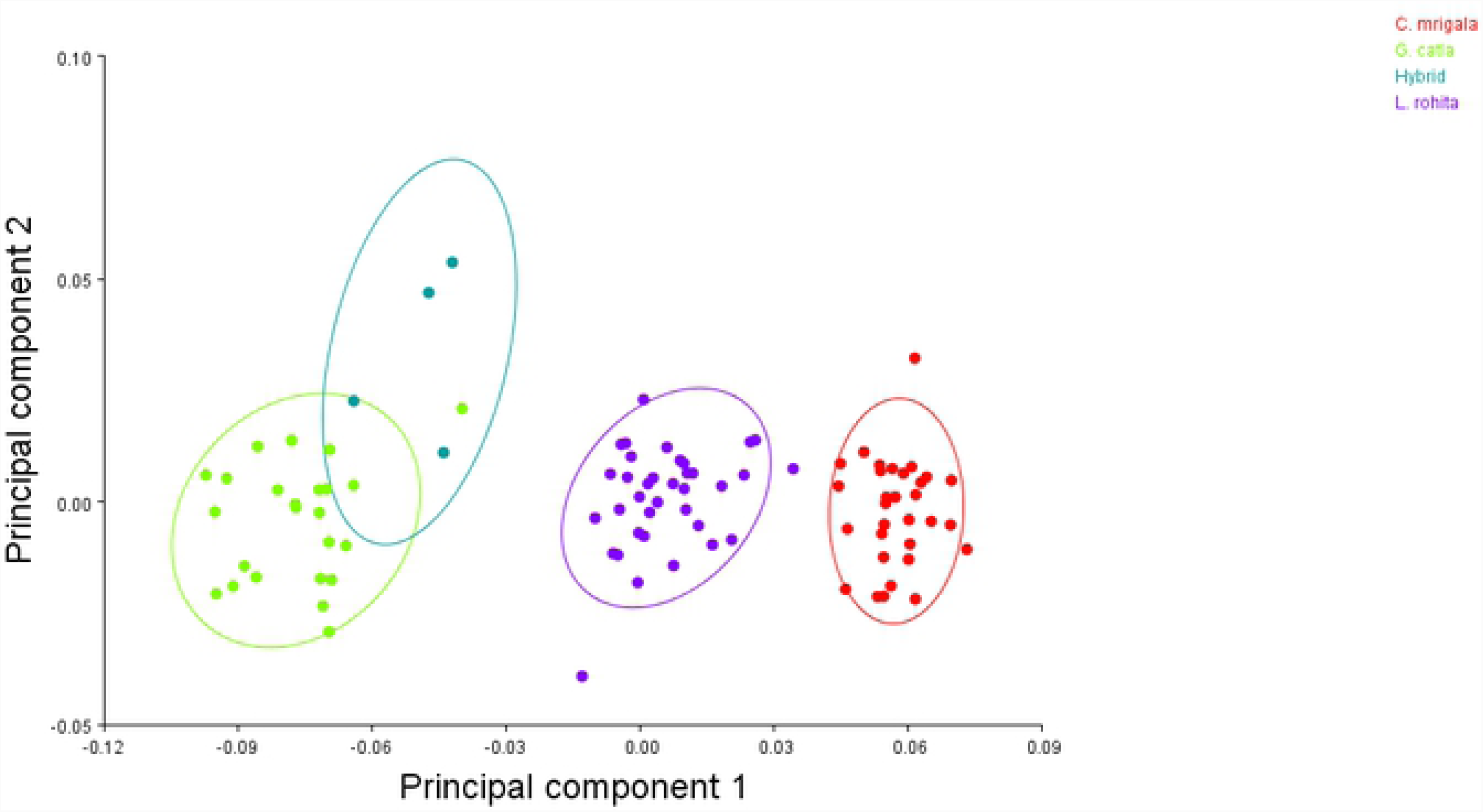
Plot of principal component analysis for Indian major carps and hybrids showing loadings of each sample on the first two principal components

**Fig. 7.**
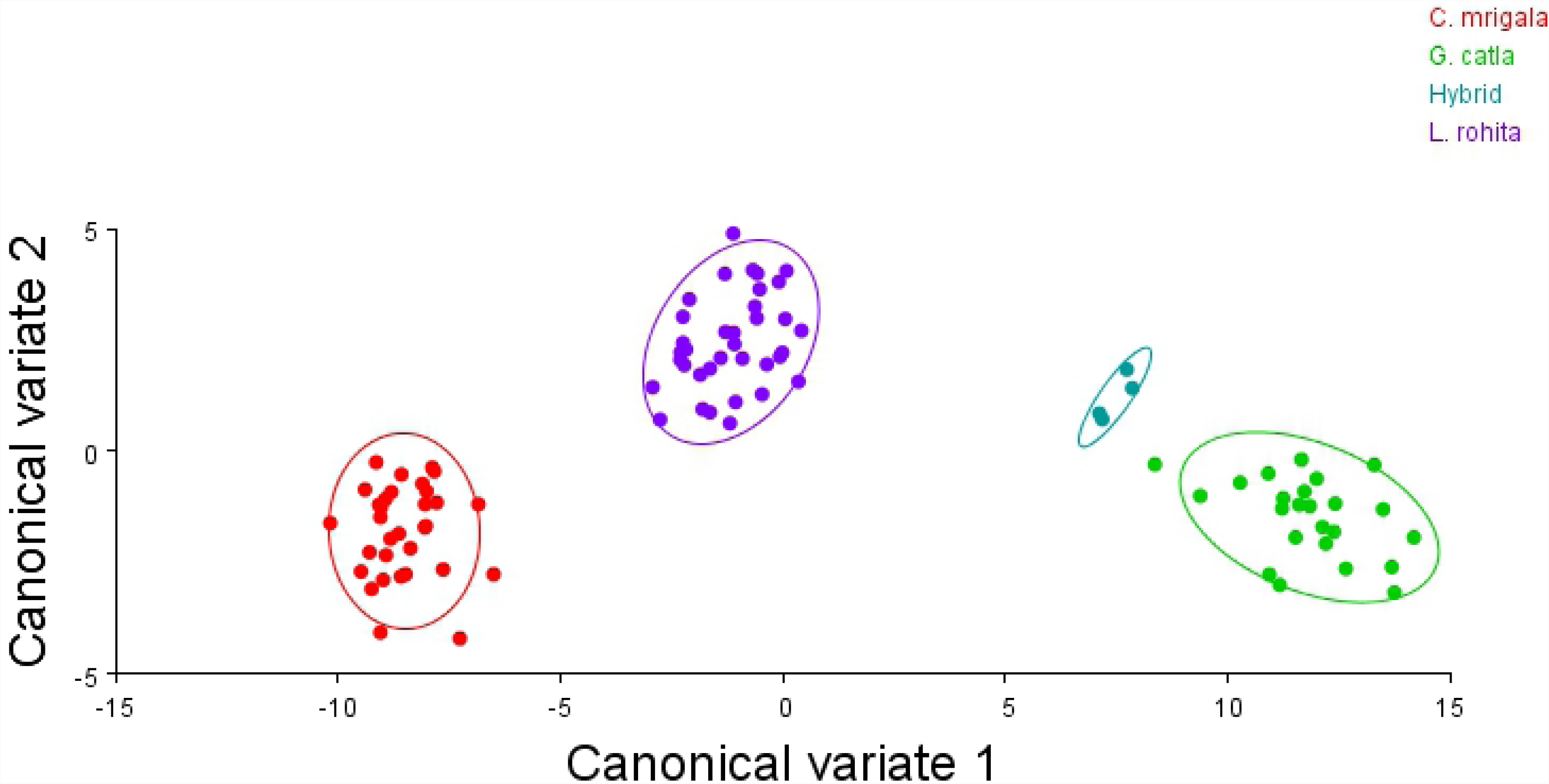
Plot of canonical variate analysis for Indian major carps and hybrids showing frequency of specimen distribution in respective group on the first two axis

## Discussion

Natural hybridization has received much attention in animal evolution in recent past [18, 21, 22, 45, 46]. Historically, majority of hybrids were assumed to be inherently unfit with reduced fertility and referred as hybrid inferiority [47, 28]. However, recent studies have provided evidence for hybrid fitness when compared to parental species [18, 45, 49-52]. Frequent interspecific hybridization encourages risk of declining natural population of indigenous fish species and thereby hamper conservation efforts [53]. Anthropogenic activities such as aquacultural activities, decline in natural population, species introductions, and loss or alteration of habitats have been frequently cited as most common contributing factors for natural hybridization [14-20, 53]. Several studies have highlighted the importance of identification, extant and causes of natural hybridization and loss of fisheries biodiversity [18, 53].

The present study focused on detecting hybrids among Indian major carps (*L. rohita, C. mrigala* and *G. catla*) collected from Ganga River along their distribution range using “Geometric Morphometric” showed four morphologically distinct groups. Size-related characters play a major job in morphometric assessment and the results may be flawed if not adjusted prior to statistical analyses of data, which were effectively removed here by Procustes analysis [37]. “Geometric Morphometric” could be a useful method to distinguish different fish species [30, 32, 33, 54, 55]. CVA showed no overlapping between groups with 100.00% correct classification in respective group. This differentiation was solely related to body depth of the fish. Samples of *G. catla* and hybrid have relatively deeper-body compared to samples of *L. rohita* and *C. mrigala*. CVA depicted hybrids position between groups of *L. rohita* and *G. catla* indicating that hybrids resultant from crossing between two species.

Geometric Morphometrics is related to change in the shape and is concerned to genetics and ecology [56]. Obtaining traditional morphometric measurements is laborious and slow process as taxonomists have to manually generate data from numerous specimens [30]. The power of landmark based morphometry to separate fish based on varying body form supports the enhancement of this technique for field based diagnosis [57]. Cadrin and Friedland [58] assessed that image processing techniques have enhanced but not been frequently applied to morphometric analyses in fisheries research.

The results obtained from the “Geometric Morphometrics” indicated that the Indian major carps showed significant phenotypic heterogeneity among the three species with clear distinctness from hybrids of *L. rohita* and *G. catla*. Cavalcanti et al [32] studied morphometric variations among six species of serranid fishes and observed non-uniform changes in body shape. Parsons et al [33] revealed high morphometric variations in two cichlid fish species using CVA. Katselis et al [59] studied morphometric variation among four grey mullet species of Mediterranean and results of DFA revealed high classification of species into their respective groups. Sakai et al [60] studied the morphological variation among three crucian carps from the Ob River system, Kazakhstan and revealed high morphometric variation among species based on landmark based morphometry. Jacquemin and Pyron [61] studied morphological variation in Cyprinidae fishes from lentic and lotic systems to gain insight into long term patterns in morphology. Reist et al [62] detected hybrids among coregonid fishes based on morphological variations. Toscano et al [18] revealed morphometric variations between the hybrids of roach (*Rutilus rutilus* L.) and bream (*Abramis brama* L.), and their parental species inhabiting an Irish lake using “Geometric Morphometrics”.

In the present study, causes of natural hybridization among Indian major carps were also recognized. It is observed that all the four individuals of hybrids were encountered upstream of barrages (Bijnor, Narora, Kanpur and Farakka) commissioned along the middle and lower stretch of Ganga River. Out of four barrages, three (Bijnor, Narora and Kanpur) are located in Uttar Pradesh and Farakka barrage in West Bengal. Starting from upstream to downstream, Bijnor barrage (1978), followed by Narora barrage (1966) and Kanpur barrage (2000) are located in middle stretch and Farraka barrage (1975) is situated in lower stretch of Ganga River.

Physical barriers commissioned along the Ganga River might have caused ecological imbalance which is evident from natural hybridization among Indian major carps. Habitat alterations are important contributing factor to the natural hybridization by facilitating geographic range expansion, creating dispersal corridors that allow movement of one species into the range of another, or reduce spawning habitat and thus limiting spawning activities [53, 63]. Physical barriers in form of barrages convert free flowing river into reservoir like habitat, affecting the ecosystem. These physical barriers restrict upward movement of fish resulting in an ecological trap for fish that ascend [64]. In the downstream area of barrages, most of the impacts are negative, while above the dam the effects are not much different [65]. Studies on loss of fish stocks by construction of physical barriers are also documented in past [65-67].

The goal of this study was to detect hybrids and their parental species among Indian major Carps (*L. rohita, C. mrigala* and *G. catla*) based on increased pace of data gathering and analysis from digital image of fish form. The results of the study showed potential of software tools in separating the parental species of Indian major carps from their hybrids using only a few body shape features, without application of DNA analysis. Therefore, this study provides a powerful tool in diagnosis of species and increasing the pace of taxonomic research.

The present study also recognizes the probable causes of natural hybridization among the Indian major carps. Habitat alteration by barrages commissioned along the distribution range of these species restricts the movement of declined fish population, thereby increasing competition with species in close proximity for spawning habitat with overlapping reproductive activities. Ranching of Indian major carps at various sites including upstream of all the barrages along the Ganga River may resolve the existing problem of natural hybridization.

## Acknowledgements

The author is thankful to the fishermen for their help in fish collection.

